# A high-fat high-sugar diet in adolescent rats impairs social memory and alters chemical markers characteristic of atypical neuroplasticity and GABAergic neurodevelopment in the medial prefrontal cortex

**DOI:** 10.1101/566026

**Authors:** Amy C Reichelt, Gabrielle D Gibson, Kirsten N Abbott, Dominic J Hare

## Abstract

Brain plasticity is a multifaceted process that is dependent on both neurons and extracellular matrix (ECM) structures, including perineuronal nets (PNNs). In the medial prefrontal cortex (mPFC) PNNs primarily surround fast-spiking parvalbumin (PV)-containing GABAergic interneurons and are central to regulation of neuroplasticity. In addition to the development of obesity, high-fat and high-sugar (HFHS) diets are also associated with alterations in brain plasticity and emotional behaviours in humans. To examine the underlying involvement of PNNs and cortical plasticity in the mPFC in diet-evoked social behaviour deficits (in this case social recognition), we exposed adolescent (postnatal days P28-P56) rats to a HFHS-supplemented diet. At P56 HFHS-fed animals and age-matched controls fed standard chow were euthanized and co-localization of PNNs with PV neurons in the prelimbic (PrL) and infralimbic (IL) and anterior cingulate (ACC) sub regions of the PFC were examined by dual fluorescence immunohistochemistry. ΔFosB expression was also assessed as a measure of chronic activity and behavioural addiction marker. Consumption of the HFHS diet reduced the number of PV+ neurons and PNNs in the infralimbic (IL) region of the mPFC by −21.9% and −16.5%, respectively. While PV+ neurons and PNNs were not significantly decreased in the ACC or PrL, the percentage of PV+ and PNN co-expressing neurons was increased in all assessed regions of the mPFC in HFHS-fed rats (+33.7% to +41.3%). This shows that the population of PV neurons remaining are those surrounded by PNNs, which may afford some protection against HFHS diet-induced mPFC-dysregulation. ΔFosB expression showed a 5-10-fold increase (*p* < 0.001) in each mPFC region, supporting the hypothesis that a HFHS diet induces mPFC dysfunction and subsequent behavioural deficits. The data presented shows a potential neurophysiological mechanism and response to specific diet-evoked social recognition deficits as a result of hypercaloric intake in adolescence.

## Introduction

The medial prefrontal cortex (mPFC) is a focal regulator of higher-order cognitive functions including working memory, cognitive flexibility, attention, social interaction and emotional regulation. The anatomy, circuity and molecular profile of the mPFC has a protracted developmental trajectory, which does not become fully refined until early adulthood.^1^ Perineuronal nets (PNNs) are extracellular matrix (ECM) assemblies that surround cortical neuronal cell bodies and proximal dendrites and are involved in the control of brain plasticity and the cessation of critical periods of development. The density of PNNs in the mPFC gradually increases from infancy through early adulthood,^2^ reflecting a key role in the closure of mPFC neurodevelopment and resulting higher-order cognitive functioning. Social cognition manifests in a similar way to working memory, but applies the principles of acquiring, storing, and manipulating information–specifically to the context of interacting in a flexible and appropriate manner with conspecifics.^3^ Abnormal social behaviour has been observed in case reports of damage to the frontal cortices in humans^4^ and rhesus monkey frontal lesion models,^5, 6^ and following mPFC disruption in rodents.^7, 8^ The developmental trajectory of the mPFC is relatively delayed compared to other cortical and subcortical regions, with its window of peak maturation across adolescence and into early adulthood.^9^

Excessive consumption of hypercaloric high-fat and high-sugar (HFHS) ‘junk food’ diets is linked not only to obesity, but also with alterations in neuroplasticity, which may manifest as deficits in cognitive and emotional processing. Junk foods are particularly appealing to children and adolescents, who are their largest consumers.

A recent body of literature examining the impact of hypercaloric diets in rodents demonstrates that the cognitive, metabolic and molecular effects of these deleterious nutritional manipulations are most prominent when consumed during the adolescent period.^10–14^

Hypercaloric diets during adolescence are also associated with neurobiological changes at the cellular and synaptic levels within the mPFC, including deficiencies in synaptic plasticity.^13^ Most notably, some of these neuronal alterations are not observed with a similar dietary exposure during adulthood, and many are more pronounced. This has given rise to the concept of critical windows of plasticity being particularly vulnerable to programming of obesity.^15^ Therefore, it is important to delineate the impact of excessive consumption of HFHS foods in adolescence on cognitive processes, and the neuronal mechanisms that underpin these alterations. Considering the late maturation of the mPFC, relative to other brain regions, it stands to reason that obesity-inducing diets during this window of neurodevelopment can have profound impacts on proper social and emotional development. Recently it has been shown that hypercaloric diet consumption in rodents can reduce parvalbumin (PV) neurons in the mPFC,^16, 17^ suggesting that concomitant alterations in primary components of brain plasticity may contribute to diet-evoked dysfunction in brain regions that control cognition and emotion. Loss of these GABAergic neurons in the mPFC are associated with cognitive deficits in diseases affecting this region, such as schizophrenia^18^ and generalised age-related cognitive decline.^19^ Considering the aforementioned evidence of hypercaloric diets causing a reduction in PV neurons, diet-induced moderate cognitive decline during mPFC maturation is clearly a potential critical window of vulnerability during proliferation of PV interneurons with key roles in mediated neurochemical development of cognitive control,^20^ memory,^21, 22^ and the generation of synaptic plasticity.^23^

A concept first proposed in 1985^24^ that is rapidly gaining traction in neuroscience is that brain plasticity is dependent not only on neurons, but also on the ECM and structures therein. The structural backbone of PNNs is hyaluronic acid (also known as a hyaluronan or hyaluronate), which is biosynthesised *via* polymerisation of glucose-derived glucuronic acid and W-acetylglucosamine by one of three hyaluronic acid synthase isoforms that are highly expressed in the rodent cortex.^25^ Structural and multifunctional chondroitin sulfate proteoglycans (CSPGs) are covalently linked to the high molecular weight hyaluronic acid biopolymer and provide an additional scaffold while directing the path of rapidly proliferating axonal connections, facilitating cell adhesion and regulating cell differentiation.^26^ In the cortex, PNNs are primarily found around fast-spiking, PV-containing GABAergic interneurons. A loss of PNNs has been observed in schizophrenia and other central nervous system diseases but the exact functional contribution of these structures or the consequences of PNN pathology are not well understood.

Of note, cortical PNNs have a delayed maturational trajectory, peaking in mid-late adolescence in rodents^27^ and humans.^2^ The closure of critical windows of plasticity is regulated by PNN expression, which increases to suppress regional plasticity.^28^ In the mature rat brain, enzymatic-induced degradation of PNNs can effectively reset PV inhibitory interneurons to a ‘juvenile state’ reflective of heightened cortical plasticity during adolescence.^29, 30^ Without PNNs present to support controlled cortical neuronal activity, firing from inhibitory PV neurons becomes erratic^31^ and the frequency of inhibitory currents from pyramidal cells decrease.^32^ These studies provide evidence that PNNs are crucial for maintaining cortical excitatory/inhibitory balance and are essential elements for the rapid and precise transmission of synaptic inhibition in the brain.

It was previously demonstrated that intermittent consumption of a HFHS diet across adolescence (postnatal days [P]28-56) reduced PV-immunopositive neurons in the infralimbic (IL) region of the mPFC in rats.^33^ In this study we examined the impact of intermittent HFHS diet on social recognition memory and associated neurobiological indicators of neuroplasticity (as PNNs and PV-interneurons) and behavioural plasticity (as ΔFosB, a biochemical marker of pathology in the natural reward pathway), and then discuss potential neurochemical mechanisms underpinning these effects.

## Experimental

### Animal ethics

All procedures and experiments were approved by the Animal Care and Ethics Committee at the University of New South Wales (Australia) and conducted in accordance with the guidelines of the Australian Code of Practice for the Care and Use of Animals for Scientific Purposes from the National Health and Medical Research Council.

### Animals and environmental conditions

Male (*n* = 24 total; 12 per group) albino Sprague Dawley rats (Animal Resources Centre, Australia) arrived in-house at 3 weeks of age (P21). Rats were randomly allocated to boxes and housed in groups of four in plastic cages with wire tops (dimensions: 60 × 40 × 26 cm) in a temperature controlled (21 ± 2 °C; humidity 55 ± 5%) colony room illuminated on a 12 h light-dark cycle (lights on 0700-1900 daily). All experiments were carried out during the light cycle between 0900 and 1700. The room in which behaviour experiments were performed was temperature (20 ± 2 °C) and humidity controlled (55 ± 5%) and illuminated to a brightness of 25 lx.

### Standard and high-fat high-sugar diets

Half of the rats were allocated to control condition (*n* = 12) or HFHS condition (*n* = 12). Standard laboratory rat chow (Gordon’s Specialty Stockfeeds, Australia) containing 65% carbohydrates, 12% fat, and 23% protein with an energy density of 11 kJ g^−1^ was available *ad libitum* throughout the experiment, as was access to normal drinking water. Rats in the HFHS diet condition were provided with 2 h daily access (between 0800 and 1000) in their home cages to semi-pure HFHS pellets (Gordon’s Speciality Stockfeeds, SP04-025; high lard modification of AIN76, semipure rodent diet). The nutritional parameters of this diet are 18.4 kJ g^−1^ digestible energy and composition of 20% fat (lard), 39.6% sucrose, and 19.4% protein, amounting to 36% energy from lipids and 55% from sucrose.

A timeline of the experimental procedures is presented in Figure 1a. Rats were given seven days from their arrival day (P21) to acclimatise to their new housing environment. During this time the animals were handled daily to habituate to the experimenters. The HFHS diet was introduced on P28 until culling at P56, coinciding with prevailing definitions of adolescence in male rats extending from P28 to ~P50-55^1, 34, 35^ that encompass the ‘periadolescent’ period when locomotor and exploratory activity are most rapidly developing.^36^ This diet was provided in addition to *ad libitum* chow and water access. Control animals did not receive any supplemented pellets. Measurements of HFHS diet consumption were calculated from the mass difference (in g) between the HFHS pellets allocated and that collected after 2 h access per cage. Body weights were recorded once at baseline before the diet began, and thereafter twice a week and averaged to calculate body weight for that week. During the diet period rats’ chow consumption over a 24 h period was recorded twice a week, in conjunction with HFHS pellet intake in the HFHS diet group and used to calculate total energy intake (in kJ). All rats in both the control and HFHS conditions were handled on week days when experimenters measured body weights and food consumption and to ensure equal habituation to experimenters.

**Fig 1.**
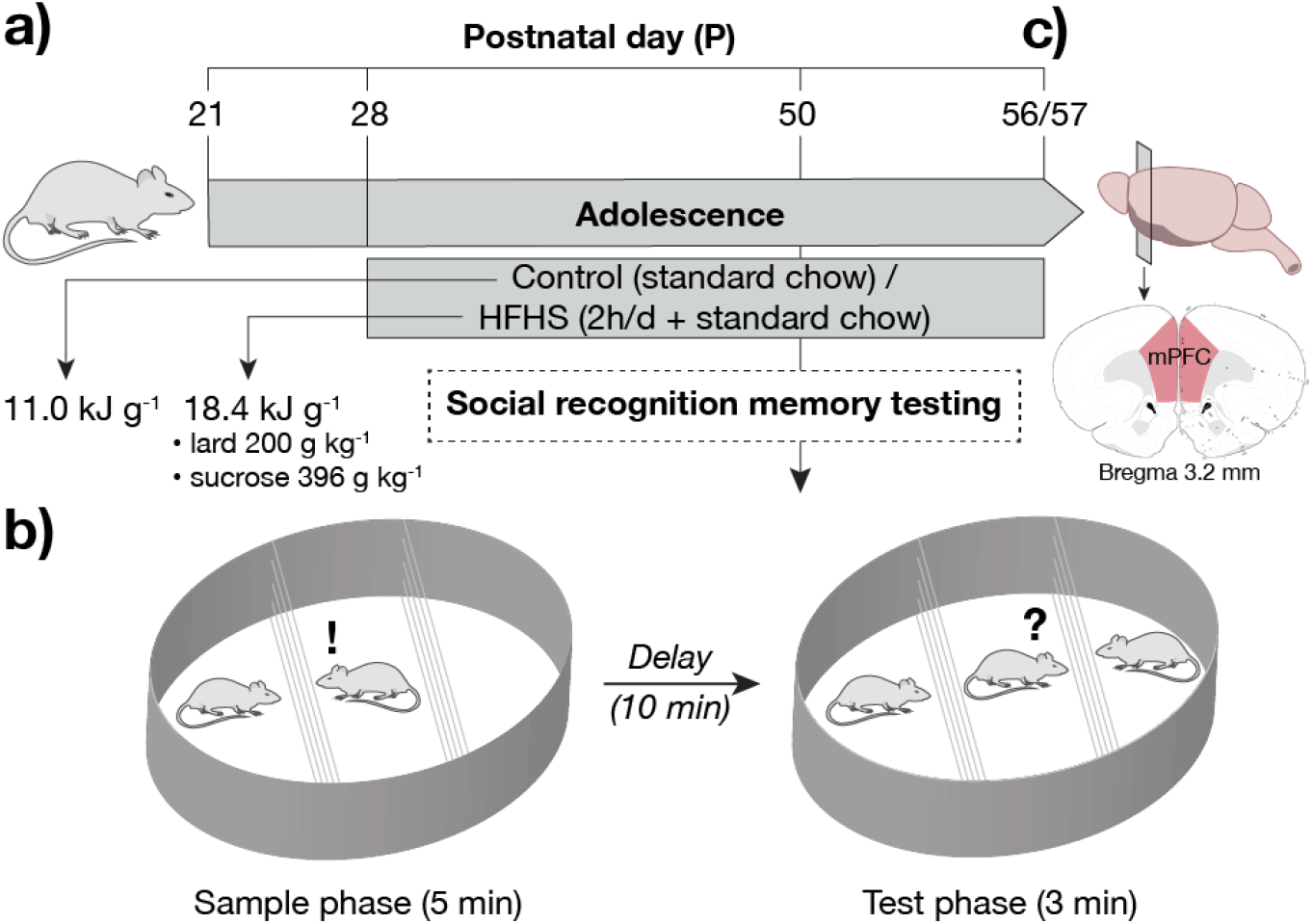
Graphical schematic of experimental design, (a) Experimental timeline. Postnatal day 21 (P21) Sprague Dawley rats were received and habituated in new housing for seven days. At P28 rats were divided into two groups (*n* = 12 per group) and fed either a diet of standard rodent chow only (total energy 11.0 kJ g^−1^) or supplemented with a HFHS diet (18.4 kJ g^−1^) containing 20.0% lard (c.a. 50% saturated and unsaturated fats) and 39.6% sucrose for two hours per day. (b) Schematic of the rat social recognition test showing the 5 min sample phase where the test rat interacted with the sample rat. At P50 animals underwent social recognition testing. Following a 10 min delay phase the test rat was reintroduced to the arena for a 3 min test phase with the sample (familiar) rat and a novel rat. c) At P56/57 the animals were euthanised and brains removed for immunohistochemical experiments. Coronal sections of the mPFC (marked in red) were taken at bregma +3.2 mm and contained regions of ACC, PrL and IL. Rat brain atlas image adapted from Swanson^37^ in accordance with a Creative Commons open access licence.

### Social memory testing

Social memory testing was based on our previously published protocol^38^ originally adapted from Crawley^39^ and Moy et al.^40^ Social memory tests were conducted in a circular arena (dimensions: 100 cm diameter, 50 cm height) constructed from grey Perspex. The arena contained two wire chambers with plastic bases (dimensions: 18 cm x 20 cm x 22 cm). The wires were interspaced 1 cm apart to allow the test rat to interact with the sample rats without physical contact. Sample, control and HFHS diet rats were habituated to the apparatus 24 h prior to testing by being placed individually into the arena with empty chambers for 10 minutes.

Social memory was tested in two phases (Fig 1b). In Phase 1, rats were placed in the arena for 5 min with one sample rat in a chamber and the other chamber left empty. Time exploring the chamber containing the sample rat *versus* the empty chamber was used as a measure of sociability. The experimental rat was then removed and placed into individual holding cages for a 5 min inter-trial interval (ITI) period. In Phase 2, the arena contained the original sample rat (familiar) in a chamber and the previously empty chamber contained a novel rat. The experimental rat was returned to the arena to explore for a 3 min period. Between test phases the arena was cleaned with 70% ethanol to eliminate odour cues.

Videos were scored to measure the duration of time the rat spent exploring the chambers during each phase. Sociability was quantified as the time spent exploring the chamber containing the sample rat as opposed to the empty chamber, and social recognition memory was measured as the time spent in proximity to the chamber containing the novel rat versus the familiar sample rat. To minimise any time effects, the order of animals tested in the social memory test was counterbalanced across groups to negate any effect of time of day when animals were tested (*i.e*. one control rat was tested, and then one HFHS rat).

### Brain extraction and sectioning

Following behavioural testing and conclusion of the 28-day diet intervention perios, rats were deeply anesthetised with sodium pentobarbital (100 mg kg^−1^, i.p.) and perfused transcardially with 0.9% saline, containing heparin (5000 i.u./mL), followed by 4% paraformaldehyde (PFA) in 0.1 M phosphate buffer (PB), pH 7.4.

Brains were then removed and post-fixed for 4 h in 4% PFA, and then placed in 20% sucrose/phosphate buffered saline (PBS) cryoprotectant solution overnight. Brains were frozen and sliced to 40 μm coronal sections on a Leica cryostat (CM1950). Four serially adjacent series of sections were obtained from each brain and stored in cryoprotectant (50% PBS, 25% ethylene glycol, 25% glycerol) at −30 °C. Sections containing the mPFC (bregma +2.2 to +3.2 mm), determined using a rat brain atlas,^41^ were stained using adjacent series for PV+PNN colocalisation, and FosB/ΔFosB immunoreactivity.

### Parvalbumin and perineuronal net immunohistochemistry

Perineuronal nets were labelled using the lectin *Wisteria floribunda* agglutinin (WFA). Although PNNs may be labelled by staining for a number of lectican-family components including aggrecan, versican, brevican, and neurocan,^42, 43^ and PNN composition does vary depending on brain region;^44, 45^ we used WFA, which binds specifically to the *N*-acetyl-D-galactosamine (GalNAc) on terminal ends of chrondroitin sulphate chains of PNNs^46^ and produces staining intensity with sufficient sensitivity for PNN counting experiments. Like all immunolabelling protocols, binding of WFA to PNNs is indirect, though it is a standard method of PNN visualisation in a variety of cortical regions.^47^ In this study we therefore refer to structures labelled by WFA as PNNs. Selective binding of WFA to GalNA β1-3 Gal is strong in the mPFC^48^ of rodents^49–51^ and humans.^2^

Free-floating sections from one series were washed in 0.1M PBS (pH 7.2) followed by blocking for 2 h at room temperature in PBS containing 5% normal horse serum (NHS), 2% normal goat serum (NGS), and 0.1% Triton X-100. Sections were incubated with fluorescein-labelled *Wisteria floribunda* lectin (1:500, FL-1351; Vector Laboratories, UK) and the primary antibody mouse anti-parvalbumin (1:2000, P3088; Sigma-Aldrich, Australia) in PBS blocking solution containing 2% NHS, 1% NGS, and 0.1% Triton X-100 for 48 h at 4 °C. After washing, sections were then incubated in Alexa Fluor 594 goat anti-mouse IgG1 (1:1000, A-21125; Life Technologies, Australia) in blocking solution for 2 h at room temperature. Sections were washed, mounted onto gelatin-coated slides and coverslipped with Vectashield+DAPI (Vector Laboratories) mounting medium. There was no immunoreactivity detected in control sections where the antibodies were omitted from the staining protocol.

### FosB/ΔFosB immunohistochemistry

Free-floating sections were washed in 0.1M PBS and blocked with 2% normal goat serum (NGS) diluted in 0.1 M PBS containing 0.1% Triton X-100. Sections were incubated at 4 °C for 48 h with a primary rabbit-raised antibody against FosB/ΔFosB (1:2000; SC-48; Santa Cruz Biotechnology, USA) diluted in PBS containing 0.1% Triton X-100 and 4% host-specific serum for 48 h at 4 °C with gentle agitation. Sections were washed and then blocked with 0.1% H_2_O_2_ before incubation with a goat anti-rabbit biotinylated secondary antibody (1:500, Vector Laboratories) in PBS containing 0.1% Triton X-100. Sections were processed by the streptavidin-horseradish peroxidase method (Vector Laboratories) and peroxidase was visualised using 3′,3′-diaminobenzadine (DAB) intensified with NiCl_2_. Sections were mounted onto 4% gelatin-coated slides, dehydrated in ascending concentrations of ethanol, cleared in histolene, and coverslipped with DPX mountant (Sigma Aldrich).

### Microscopy and immunoreactivity analysis

All slides were coded and imaged/analysed blind to treatment. Fluorescent and brightfield images were captured on an Olympus BX53 microscope (Olympus, Australia) equipped with a digital camera (DP72) and a fluorescent lamp (X-Cite, Series 120Q). The prelimbic (PrL), infralimbic (IL), and anterior cingulate (ACC) subregions of the mPFC were counted from 3 consecutive sections from the left hemisphere between bregma +3.2 and +2.7 mm.

Regions were delineated using clearly visible landmarks and predefined boundaries according to a rat brain atlas.^41^ When landmarks did not clearly delineate regions, images were cropped to boundaries created with a 10 × 10-grid reticule located in the left eye piece of the microscope using a 10× objective lens. The eye-piece grid was aligned with the dorsal aspect of the forceps minor corpus callosum to define the top of the PrL and base of the ACC, and outlined boundaries of a 1 mm (high) region between the medial edge of the section and the forceps minor (PrL) and a 0.7 mm high region for the ACC. For the IL a 0.7 mm high region was created with the grid aligned at the point where the forceps minor and the edge of the section become parallel. Total counts of PV-IF neurons and PNNs were manually counted using ImageJ (v1.46)^52^ using a 10× objective. FosB/ΔFosB-immunoreactive cells were counted automatically using CellSens (Olympus) software within 1 mm × 1 mm (PrL) or 1 mm * 0.7 mm (IL and ACC) region areas. Cells were considered positive for FosB/ΔFosB if the stained nucleus was the appropriate size (area range = 10-40 μm^2^) and shape (at least 50% of circularity) and was distinct from the background (exceeding a threshold of −12 to −255 [black]).

### Scoring, exclusion criteria, and statistical analysis

An observer blind to the group allocations manually scored video recorded trials of the social recognition memory test using Macropod ODlog^®^ software. Statistical analyses were conducted using SPSS Statistics 23 (IBM, USA) and Prism v8.0 (GraphPad, USA). Body weight and energy intake data were analysed using a mixed-design ANOVA or repeated measures ANOVA for energy consumption within the HFHS group. Simple main effects explored interactions in more detail. An alpha value was set at 0.05.

## Results

### Dietary effects on weight gain and energy intake

Control and HFHS-diet rats both gained weight across the 28-day diet manipulation period (*F_(9,99)_* = 1002, *p* < 0.001). Consistent with our previous observations where significant differences in mean body weight between control and HFHS-diet rats only become apparent after >4 weeks of diet manipulation,^17^ weights between groups did not significantly differ during the specific window representing adolescence used (main effect diet: *F* < 1, diet x time: *F_(9,99)_* = 1.84, *p* = 0.07). Hypercaloric diets during this 28-day period have a well-established phenotype of impaired cognition irrespective of body mass,^16^ suggesting subtle neurological effects precede measurable changes in body mass. Over the 28-day diet manipulation period, rats in both groups significantly increased their energy consumption (*F_(3,6)_* = 20.8, *p* < 0.01), though the overall energy intake did not differ between control and HFHS-diet rodents (main effect diet: *F_(1,2)_* = 3.93, *p* = 0.19, diet x time: *F_(3,6)_* = 2.01, *p* = 0.21).

**Fig 2.**
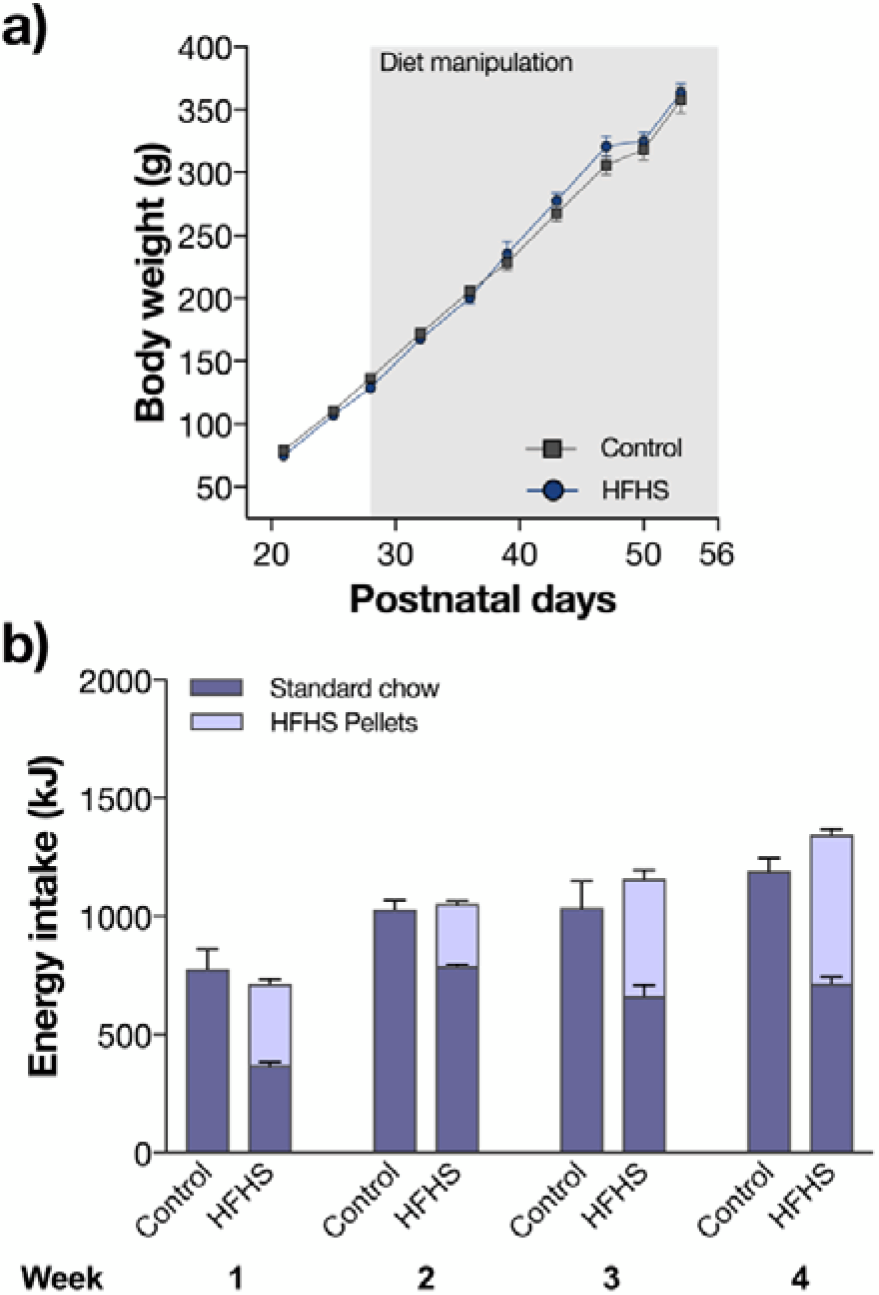
Effects of HFHS diets on body mass and energy intake. a) Mean body weights of control and HFHS rats across the study. b) Mean energy consumption (kJ) per cage of rats across the 4-week diet exposure period. Error bars represent +SEM.

### High-fat high-sugar diet impairs social recognition memory

Social behaviour has been typically examined in mice using the ‘three-chamber’ social approach test. We have successfully adapted this protocol for use in rats to examine whether changes in social recognition memory.^38^ During the ‘sample’ sociability phase of the test, both control and HFHS rats preferentially explored the novel, or ‘sample’ rat compared to the empty cage (*F_(1,11)_* = 688.2, *p* < 0.001) with no significant between group (*F* < 1) or interaction effect (*F* < 1; Fig. 3a). However, social recognition was impaired in HFHS rats in the ‘test’ phase, which explored the familiar and novel rat equally, contrasting to the strong preference of control rats to explore and interact with the novel rat (chamber x diet group interaction, *F_(1,11)_* = 84.2, *p* < 0.001; control *F_(1,11)_* = 174.5, *p* < 0.001, HFHS *F_<1_*; Fig. 3b).

**Fig 3.**
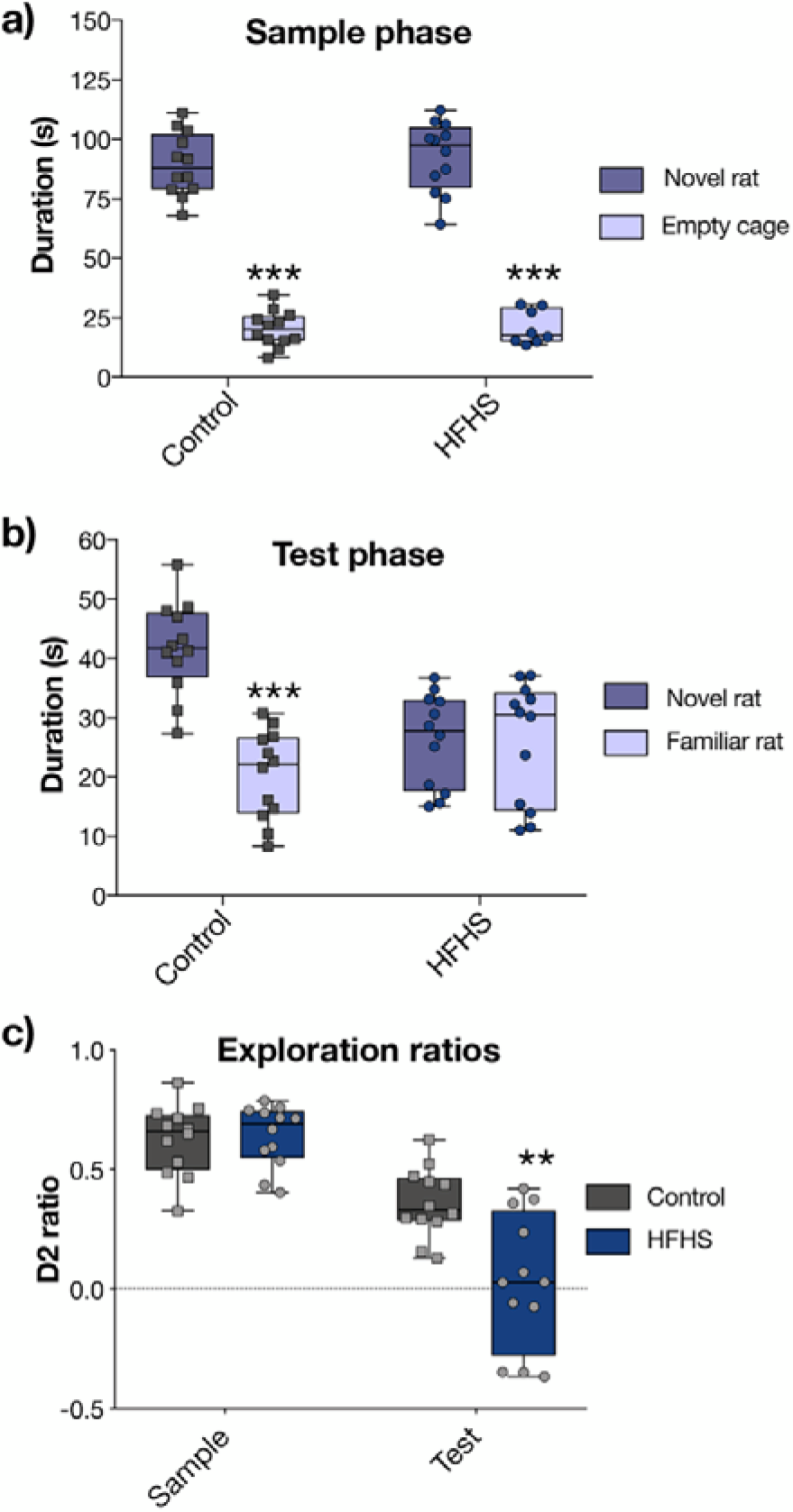
A HFHS diet impairs social memory. a) Both control and HFHS-diet rats preferentially interacted with the novel rat than empty cage during sociability testing (*** *p* < 0.001 *vs* empty chamber). b) In the test phase, only controls showed a preference to the novel rat over the familiar animal (*p* < 0.001), with HFHS-diet rats showing no statistically-significant difference in social interaction with the novel animal (*p* = 0.83). c) This was also reflected in the significant decrease in D2 ratio (time novel – time familiar) / total time exploring) between groups during the test phase only (** *p* < 0.01).

Exploration ratios were statistically compared using the D2 scoring method (*t_novel_ – t_familiar_*) / (*t_novel_ + t_Familiar_*)^53, 54^ for the sample and test phases. The D2 ratio can vary between +1 and −1, with a positive ratio showing a preference for novelty, and a score of 0 indicating equal exploration. During the sample phase, a strong preference for the cage containing the rat over the empty chamber was observed in both control (mean ± SEM: control = 0.63 ± 0.04) and HFHS rats (0.64 ± 0.04), with no significant difference between diet groups (*F_(1,11)_* = 1.31, *t_(2,22)_* = 0.27, *p* = 0.79; Fig 3c). The D2 ratio differed significantly between diet groups in the test phase only (0.36 ± 0.04; HFHS = 0.03 ± 0.08; *F_(1,11)_* = 3.78, *t_(2,22)_* = 3.62, *p* < 0.01).

### Effects of a hypercaloric diet on GABAergic interneuron and perineuronal net numbers in the medial prefrontal cortex

Parvalbumin neurons were identified in all cortical layers with the exception of layer I; a distribution that is consistent with previous reports.^55^ The PNNs appeared mostly in the mid-to deep-cortical layers (*e.g*. IV and V/VI), as was found in previous studies with adult rats.^32, 50^

Perineuronal nets preferentially enshroud PV interneurons during late adolescence,^56^ though consistent with other reports^57^ PV +ve neurons were both surrounded by and present without an associated PNN (Fig 4a). Across the mPFC, HFHS diet consumption significantly reduced the total numbers of PV +ve neurons (mean ± SEM; control = 134.3 ± 7.3, HFHS = 101.2 ± 10.7; *t_(2,22)_* = 2.55, *p* < 0.05) and PNNs (control = 160.9 ± 5.8, HFHS = 143.4 ± 5.4; *t_(2,22)_* = 2.22, *p* < 0.05). In the three regions analysed separately, there was a trend to lower numbers of PV +ve neurons in the PrL (*t_(2,22)_* =1.156, *p* = 0.25), though this only reached significance in the ACC (*t_(2,22)_* = 2.28, *p* < 0.05) and IL (*t_(2,22)_* = 2.30, *p* < 0.05; Fig. 4b) in HFHS diet rats. This decrease in PV interneurons in the IL of HFHS diet rats corresponded with a decrease in the total number of PNNs showing immunofluorescence after fluorescein-conjugated *Wisteria floribunda* lectin labelling (*t_(2,22)_* = 2.29, *p* < 0.05; Fig. 4c), however this reduction was not significant in the ACC (*t_(2,22)_* = 1.67, *p* = 0.10) or PrL (*t* < 1). Notably, there was a significant increase in the percentage of PV interneurons co-localised with PNNs in all three subregions analysed in the HFHS-diet group (ACC *t_(2,22)_* = 4.97, *p* < 0.001; PrL *t_(2,22)_* = 3.53, *p* < 0.001; IL *t_(2,22)_* = 3.90, *p* < 0.001; Fig. 4d-f), beyond that expected of normal adolescent rats.^56^

**Fig 4.**
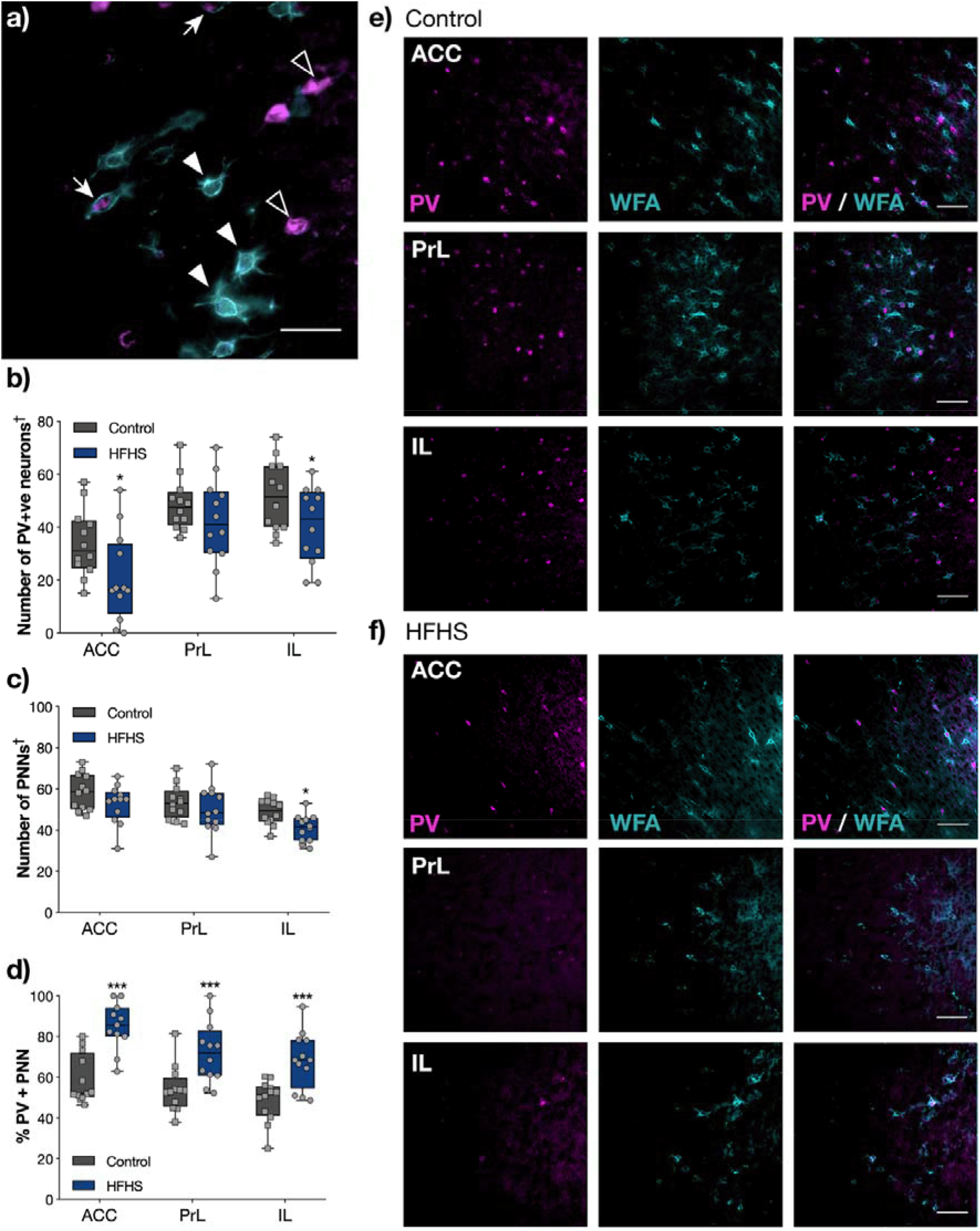
A HFHS diet alters PV and PNN numbers in the mPFC. a) PNNs (labelled with fluorescein-conjugated *Wisteria floribunda* lectin [WFA]; cyan) were both associated with PV +ve neurons (magenta) and other non-PV-containing neurons, and PV +ve neurons were also found absent of a surrounding PNN in the control IL region of the mPFC at P56/57. Scale bar = 20 μm. b) PV +ve neurons were decreased in the ACC and IL (p < 0.05) and c) the number of PNNs were significantly decreased (* *p* < 0.05) in the IL region of HFHS-diet rats compared to controls. † denotes units in mm^2^; ACC = 0.7, PrL = 1, IL = 0.7. d) In the ACC, PrL and IL regions there was a marked increase (\*\*\**p* < 0.001) in the percentage of PV interneurons with PNNs in the HFHS-diet group *vs* control group. e-f) Dual immunofluorescence images from each subregion confirmed both a generalised decrease in PV neurons, PNNs, and increased number of PNN-associated PV +ve cells in HFHS-diet rats compared to control. Scale bar = 50 μm.

### FosB/ΔFosB immunoreactivity in the medial prefrontal cortex of high-fat high-sugar fed rats

FosB is a key regulator of cell proliferation, differentiation, and transformation; and is thus an important marker of neuroplasticity.^58^ ΔFosB is a splice variant of the *Fosb* gene that is highly stable and plays a major role in long-term adaptation behaviours,^59^ including modified processing of natural rewards and associated modifications to behaviours.^60^ FosB/ΔFosB immunoreactivity (Fig 5a) was significantly increased in all three regions of the mPFC in HFHS diet rats (ACC *t_(2,22)_* = 6.66, *p* < 0.001; PrL *t_(2,22)_* = 5.09, *p* < 0.001; IL *t_(2,22)_* = 2.75, *p* < 0.01; Fig 5b) indicating a marked long-term transcriptional effect on a protein linked to addiction-like behaviours including drug-seeking^61^ and overconsumption of food;^62^ and social impairment in rodents.^63^

**Fig 5.**
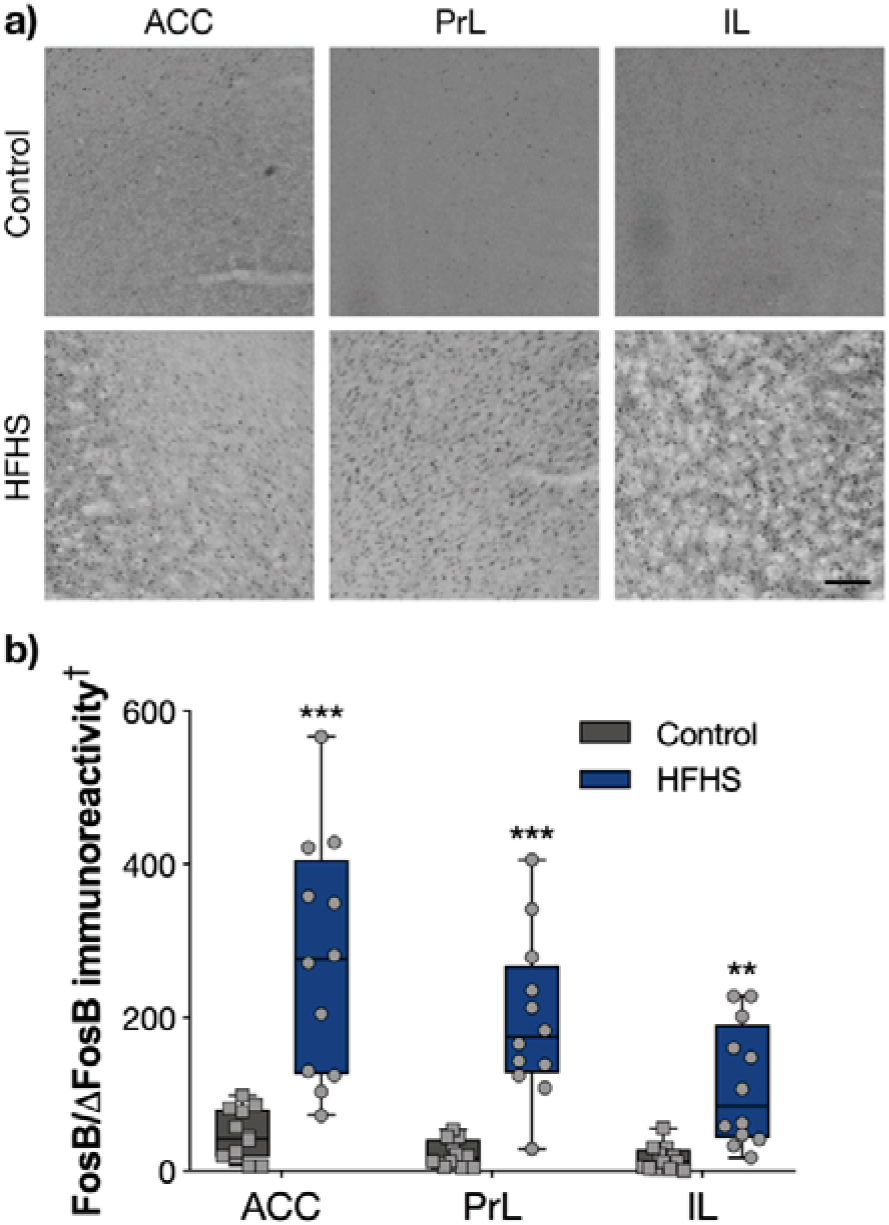
A key marker of neuroplasticity is altered by HFHS diet intake during adolescence. a) FosB/ΔFosB immunoreactivity was increased in the three mPFC subregions analysed (scale bar = 100 μm), b) showing an approximately five-fold increase in the ACC, PrL (*** *p* < 0.001), and IL (** *p* < 0.01) compared to the control group. † denotes units in mm^2^; ACC = 0.7, PrL = 1, IL = 0.7.

## Discussion

Parvalbumin-expressing neurons mature across early life. In rodents, PV positive cells typically first appear in the sensory cortices by the second postnatal week (P8-11), and in associative cortices by P11-13, with adult patterns of expression in the cortex being reached by the juvenile stage of development (*i.e*. P24/P25).^64^ Similar PV neuron counts are observed in the PrL of juvenile (P25) and adolescent (P40) rats,^65^ PrL and IL of adolescent (P35) and adult (P70) rats,^56^ and also the ACC in juvenile (P20), adolescent (P40), and adult (P90) mice.^66, 67^

Our findings shows that daily intermittent consumption of a HFHS diet during adolescence leads to behavioural deficits in social recognition memory, a task that has been directly linked to mPFC function.^7^ Moreover, consumption of this diet significantly reduced numbers of PV im mu noreactive neurons in the ACC and IL regions of the mPFC, as well as decreased PNNs in the IL. Neural circuitry in the ACC is includes input from the social behaviour-regulating limbic system and cognitive functions of the mPFC,^68^ while the IL is a region critical for behavioural regulation and cognitive flexibility.^69^ We also observed an increase in the percentage of PV immunoreactive neurons that were surrounded by PNNs in rats that consumed a HFHS compared to controls. This indicates that the majority of PV neurons in HFHS diet fed animals are those surrounded by PNNs, which may afford some protection against HFHS diet-induced mPFC-dysregulation.^70^ Previous studies with both rats and mice fed hypercaloric diets across the adolescent period have demonstrated impairments in the extinction of learned fear behaviour,^13, 17^ which has been shown to depend on IL cortex integrity.^71, 72^ Similar diets have been shown to evoke behavioural abnormalities following adolescent diet manipulation at the molecular levels, including deficiency of the ECM glycoprotein reelin^13^ and evidence of transcriptional reprogramming in 38 individual mRNAs.^73^ We have observed shortterm memory deficits in rats on hypercaloric diets previously,^38, 74, 75^, and further understanding of the effects HFHS diets have on social memory and associated deficiencies using this experimental paradigm would require detailed analysis of the hippocampus and how it related to mPFC dysfunction.

Considering that GABAergic neurotransmission function switches from excitatory to inhibitory during maturation,^76^ dysfunction or loss of mPFC PV interneurons is likely to evoke a shift in the cortical excitatory/inhibitory (EI/) balance^77^ and resulting cognitive phenotypes.^66, 78-80^ Data suggests that E/I balance helps facilitate the induction of long-term potentiation, which could be critical in allowing the mPFC to remain flexible and adapt to the environment^81, 82^ and in the performance of working memory tasks.^8^ This supports our observation of increased FosB/ΔFosB immunoreactivity across the ACC, PrL and IL regions of the mPFC in HFHS-fed rats, indicating that HFHS diet has an impact on chronic neural activity within the mPFC. This also supports a previous observation of significantly increased FosB immunoreactivity in the IL in rats following 7 weeks of intermittent HFHS, however we saw increases across all regions of the mPFC, indicating that there may be elements of compensation in neural activity levels after adolescence.^17^

Recent studies show that PNNs—specialised ECM structures that form around the synapses on the cell soma and proximal dendrites of primarily fast-spiking, PV-containing GABAergic interneurons^83^ are critical for the control of central nervous system plasticity, maintenance and repair.^84, 85^ The enzymatic degradation of chondroitin sulfate proteoglycans (CSPGs)—major components of PNNs—can reopen the juvenile critical period of heightened cortical plasticity.^30^ Similarly, enhanced neuroplasticity is observed in adult mice lacking the *Crtl1/Hapln1* gene, which encodes a link protein essential for formation of PNNs, and ablation of this gene leads to impaired CSPG organisation and PNNs with decreased structural integrity.^86^ Removal of PNNs *via* enzymatic degradation with chondroitinase-ABC has been shown to enhance excitatory synaptic transmission and facilitate long-term depression, suggesting that PNNs may be critical in the regulation of GABA-mediated inhibition.^87, 88^ This is consistent with findings that a high proportion of parvalbumin-containing GABAergic (PV) interneurons are covered with PNNs in healthy cortex.^51^

Highly active neurons are often found surrounded by PNNs, and it has been proposed that they provide a buffered, negatively charged environment that supports neuronal activity.^51^ Perineuronal nets are thought not only to regulate neuroplasticity, but also to also protect neurons from oxidative stress caused by neuroinflammation,^67^ providing a strong rationale for their involvement in HFHS-diet induced cognitive decline, even over short-term periods.^89^ Dietary-induced obesity is known to evoke neuroinflammation in the brain,^90^ and while most attention has been paid to its role in promoting oxidative stress in chronic neurodegenerative disorders or acute head injury, a sustained elevation in oxidative stress may cause more subtle neurochemical changes, such as those observed on the dynamic PV/PNN network in the mPFC of HFHS-diet adolescent rats. Indeed, an obesity-inducing diet in aged mice showed a marked increase in neuroinflammation and markers of oxidative damage compared to younger adult controls.^91^ Moreover, PV neurons without a surrounding PNN have been shown to be highly vulnerable to oxidative stress, particularly during key windows of development,^92^ providing a biochemical rationale for why consumption of HFHS diets have a more pronounced detrimental impact during adolescence when PNNs are yet to fully form around PV neurons in the mPFC.^17^ Further research using this experimental paradigm should involve investigation of oxidative stress markers, such as total protein carbonyl levels of immunostaining of 8-hydroxyguanosine and derivatives.

If increased neuroinflammation and oxidative stress are indeed the underlying chemical cause by which a HFHS diet causes disruption to mPFC neuroplasticity and function, there are several candidate mechanisms that may be involved and are worthy of further study. The volume of research into oxidative stress and schizophrenia, which affects the same neuroanatomical regions as those examined here, may provide some clues into the major chemical effects of HFHS-induced inflammation and production of reactive oxygen species (ROS) that were comprehensively reviewed by Bitanihirwe and Woo (2011).^93^ Of particular interest to us is the effect on neurotransmitter metabolism, considering its indisputable links to cognitive development in the mPFC. GABA produced by PV interneurons does not directly interact with ROS, though oxidative stress does reduce activity of GABA_A_-gated chloride channels,^94^ potentially impairing regulation of plasticity in the mPFC. Indeed, direct determination of GABA levels in the frontal cortex of adult rats fed a hypercaloric diet by liquid chromatography showed an approximately 50% in the treated animal.^95^ Other neurotransmitters present in the mPFC may also be adversely affected by increased ROS generation following a HFHS-diet; glutamate neurotoxicity has an increasing body of evidence suggesting a potentiating role for oxidative stress,^96^ while autooxidation of dopamine to potent neurotoxic quinones is promoted by excess redox-active iron^97^ present as a result of neuroinflammation, or a potential increase in pathological levels of ferroptosis–a newly-discovered iron-dependent, non-apoptotic cell death pathway found in multiple disorders sharing underlying neurochemical features with hypercaloric diets.^98^ Development of accurate and precise neurotransmitter metabolomic methods that allow simultaneous and quantitative detection of neurotoxic metabolites potentially involved in driving oxidative stress in the mPFC are crucial for future studies examining the relationship between neurotransmitters, their impaired role in regulating neuroplasticity, and possible contributory mechanisms to neuropathology.

## Conclusions

In this study, we observed functional changes to the mPFC of rats exposed to a HFHS diet across the adolescent period, which manifested behaviourally as impaired social recognition. Moreover, dual immunofluorescence revealed that the HFHS diet reduced PV neurons, and altered the co-expression of PNNs, elements of the extracellular matrix with critical involvement in the regulation of neuroplasticity and protection of developing neurons. In particular, the increased percentage of PV neurons that co-expressed PNNs leads to the conclusion that this population of neurons were those that are protected from HFHS-diet induced neurochemical changes, however further studies are required to fully determine the underlying mechanisms for this alteration in neuronal populations

## Acknowledgements

This work was supported by an Australian Research Council Discovery Early Career Research Award (DE140101071) and Discovery Project (DP180101974) to ACR. DJH was supported by a National Health and Medical Research Council-administered Career Development (Industry) Fellowship (GNT1122981). The Florey Institute of Neuroscience and Mental Health acknowledges the strong support from the from the Victorian Government and in particular the funding from the Operational Infrastructure Support Grant.

